# HTSlib - C library for reading/writing high-throughput sequencing data

**DOI:** 10.1101/2020.12.16.423064

**Authors:** James K. Bonfield, John Marshall, Petr Danecek, Heng Li, Valeriu Ohan, Andrew Whitwham, Thomas Keane, Robert M. Davies

## Abstract

**Abstract:** *Background:* Since the original publication of the VCF and SAM formats, an explosion of software tools have been created to process these data files. To facilitate this a library was produced out of the original SAMtools implementation, with a focus on performance and robustness. The file formats themselves have become international standards under the jurisdiction of the Global Alliance for Genomics and Health.

*Findings:* We present a software library for providing programmatic access to sequencing alignment and variant formats. It was born out of the widely used SAMtools and BCFtools applications. Considerable improvements have been made to the original code plus many new features including newer access protocols, the addition of the CRAM file format, better indexing and iterators, and better use of threading.

*Conclusion:* Since the original Samtools release, performance has been considerably improved, with a BAM read-write loop running 5 times faster and BAM to SAM conversion 13 times faster (both using 16 threads, compared to Samtools 0.1.19). Widespread adoption has seen HTSlib downloaded over a million times from GitHub and conda. The C library has been used directly by an estimated 900 GitHub projects and has been incorporated into Perl, Python, Rust and R, significantly expanding the number of uses via other languages. HTSlib is open source and is freely available from htslib.org under MIT / BSD license.

*Contact:*

## Background

When the 1000 Genomes Project [1] was launched in early 2008, there were many short-read aligners and variant callers. Each of them had its own input or output format for limited use cases and interoperability was a major challenge. Users were forced to implement bespoke format converters to bridge tools, and since formats encoded different information, this was time consuming, laborious and sometimes impossible. This fragmented ecosystem hampered the collaboration between the participants of the project and delayed the development of advanced data analysis algorithms.

In a conference call on 21st October, 2008, the 1000 Genomes Project analysis subgroup decided to take on the issue by unifying a variety of short-read alignment formats into the Sequence Alignment/Map format or SAM for short. Towards the end of 2008 the subgroup announced the first SAM specification, detailing a text-based SAM format and its binary representation, the BAM format [2]. SAM/BAM quickly replaced all the other short-read alignment formats and became the *de facto* standard in the analysis of high-throughput sequence data. In 2010, a Variant Call Format (VCF) was introduced for storing genetic variation [3]. Later, in 2011, as the number of sequenced samples grew and the text format proved too slow to parse, a binary version BCF [see Additional file, Figure S1] was developed[4].

The SAM/BAM format originally came with a reference implementation, SAMtools [2], and VCF/BCF with VCFtools [3] and BCFtools (then part of the SAMtools package). Numerous other tools have been developed since then in a wide variety of programming languages. For example, HTSJDK is the Java equivalent [5] and is used extensively in Java applications; Sambamba [6] is written in the D language and focuses primarily on efficient multi-threaded work; Scramble [7] has BAM and SAM capability and is the primary source for experimental CRAM [8,9] development; and JBrowse [10] implements read-only support for multiple formats in JavaScript.

While the original implementation of SAMtools and BCFtools provided application programming interfaces (APIs) to parse the files, it mixed these APIs with application code. This did not guarantee long-term stability and made it difficult to interface in other programs. To solve this, in 2013 the decision was taken to separate the API from the command line tools and to produce HTSlib as a dedicated programming library that processes common data formats used in high-throughput sequencing. Support for the European Bioinformatics Institute’s CRAM format was added and in 2014 the first official release (1.0) was made. This library implements stable and robust APIs that other programs can rely on. It enables efficient access to SAM/BAM/CrAM, VCF/BCF, FASTA, FASTQ, block-gzip compressed data and indexes. It can be used natively in C/C++ code and has bindings to many other popular programming languages, such as Python, Rust and R, boosting the development of sequence analysis tools.

HTSlib is not merely a separation; it also brought numerous improvements to SAMtools, BCFtools and other third-party programs depending on it. HTSlib is linked into approximately 900 GitHub projects [see Additional file, Section S2], and HTSlib itself has been forked more than 300 times. HTSlib has been installed via bioconda over 1 million times, and there are around 10,000 GitHub projects using it via Pysam. The library is freely available for commercial and non-commercial use (the MIT / BSD compatible license) from htslib.org and GitHub.

## Findings

### Implementation

The main purpose of HTSlib is to provide access to genomic information files, both alignment data (SAM, BAM and CRAM formats) and variant data (VCF and BCF formats). The library also provides interfaces to access and index genome reference data in FASTA format and tab-delimited files with genomic coordinates.

Given the typical file sizes of genomic data, compression is necessary for efficient storage of data. HTSlib supports a GZIP-compatible format BGZF (Blocked GNU Zip Format) which limits the size of compressed blocks, thus allowing indexing and random access to the compressed files. HTSlib includes two standalone programs that work with BGZF; *bgzip* is a general purpose compression tool while *tabix* works on tab delimited genome coordinate files (e.g. BED and GFF) and provides indexing and random access. BGZF compression is also used for BAM, BCF and compressed FASTA files. The CRAM format uses column-specific compression methods including gzip, rANS[11], LZMA and bzip2. The CRAM implementation in HTSlib learns the best performing compression method on the fly [see Additional file, Section S3].

The HTSlib library is structured as follows: the media access layer (Figure 1a) is a collection of low-level system and library *(libcurl, knet)* functions, which facilitate access to files on different storage environments (disk, memory, network) and over multiple protocols to various online storage providers (AWS S3, Google Cloud, GA4GH htsget [12]; Figure 1b). This functionality is transparently available through a unified low-level stream interface hFILE (Figure 1c). All file formats (Figure 1d-e) are accessible through a higher-level file-format agnostic htsFILE interface, which provides functions to detect file types, set write options and provides common code for file iterators. Building on this layer are specialisations for alignment (SAM, BAM and CRAM) and variant (VCF and BCF) files and various auxiliary functions (Figure 1f-g).

**Figure 1:**
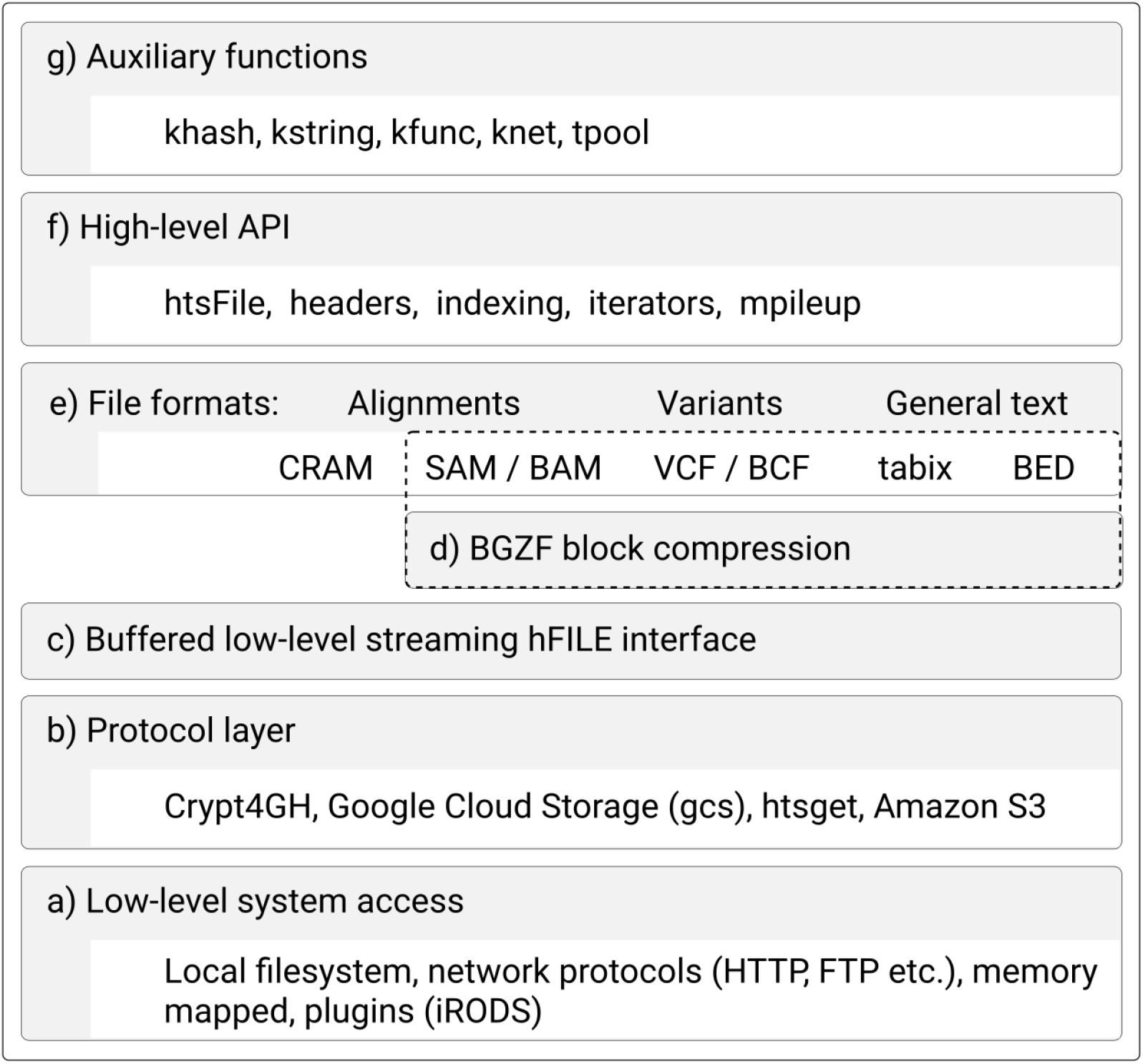
Htslib design.

This API (Figure 1f-g) can be roughly divided into several classes:

1. The **File Access API** has basic methods for opening and closing alignment and variant files, as well as reading and writing records to a file. HTSlib automatically determines the input file type by its contents and output type by filename. Further explicit control is provided for format, data layout (in CRAM) and file compression levels. Data sorted in genomic order may also be indexed at the time of writing (for alignment data) or at a later post-writing stage.
2. The **Header API** is a collection of methods that enables extensive control of SAM and VCF headers, including reading, writing and parsing the header, accessing and updating individual tags, adding and removing header lines.
3. The **Data API** provides methods for parsing, updating and retrieving information from individual record fields on both alignment and variant data. The library also includes the ability to read multiple VCF and BCF files in parallel, transparently merging their contents, so that the reader can easily process records with matching genomic positions and alleles.
4. The **Index / Iterator API** offers the ability to extract information from the various index formats specific to genomic data (BAI, CSI, CRAI, TBI), and to create iterators for genomic files. The original BAI and TBI indices were limited to 512 Mbases and were replaced by CSI allowing up to 2^44^ bases. Both sequence alignment and variant call formats have millions of records which can be indexed by genomic location. An iterator groups a list of target genomic regions into a list of file offsets and contains the stepping and filtering logic to allow the file reader to extract only the information of interest. Additionally, the library provides the *regidx* API which allows to efficiently search and intersect regions from arbitrary row-oriented text formats.
5. The **Mpileup API** performs a data pivot. Alignment data in SAM, BAM and CRAM is retrieved in row-oriented format, record by record. Data rotation (merging one or more input files) presents the sequence data in a column-oriented form per reference position. This information can be used for SNP and indel calling, consensus generation and to make alignment viewers. Mpileup can also optionally calculate base alignment quality scores (BAQ) for each read [13]. The BAQ scores can be used to reduce false-positive SNP calls by lowering the confidence scores at locations where the read alignment may be incorrect.
6. HTSlib also includes various utility convenience functions such as hash tables, string manipulation, linked lists, heaps, sorting, logging, and ensures portability between big- and little-endian platforms. Many of these originate from Klib[14]. A thread pool interface is provided for general multi-threading.

### Benchmarks of sequence alignment formats

Given the widespread use of the library, performance and low memory requirements are paramount, which means even relatively small improvements can lead to time and energy savings when analyzing large amounts of data.

To test maximum throughput for alignment data, elapsed times were obtained for each file type using both 1 main thread and also 16 additional worker threads. Not all tools supported indexing of all formats, and only in more recent HTSlib versions is there support for indexing and random access of BGZF compressed SAM files. Full benchmarks are in Additional File, Table S5, with a summary for BAM shown here in Figure 2.

**Figure 2:**
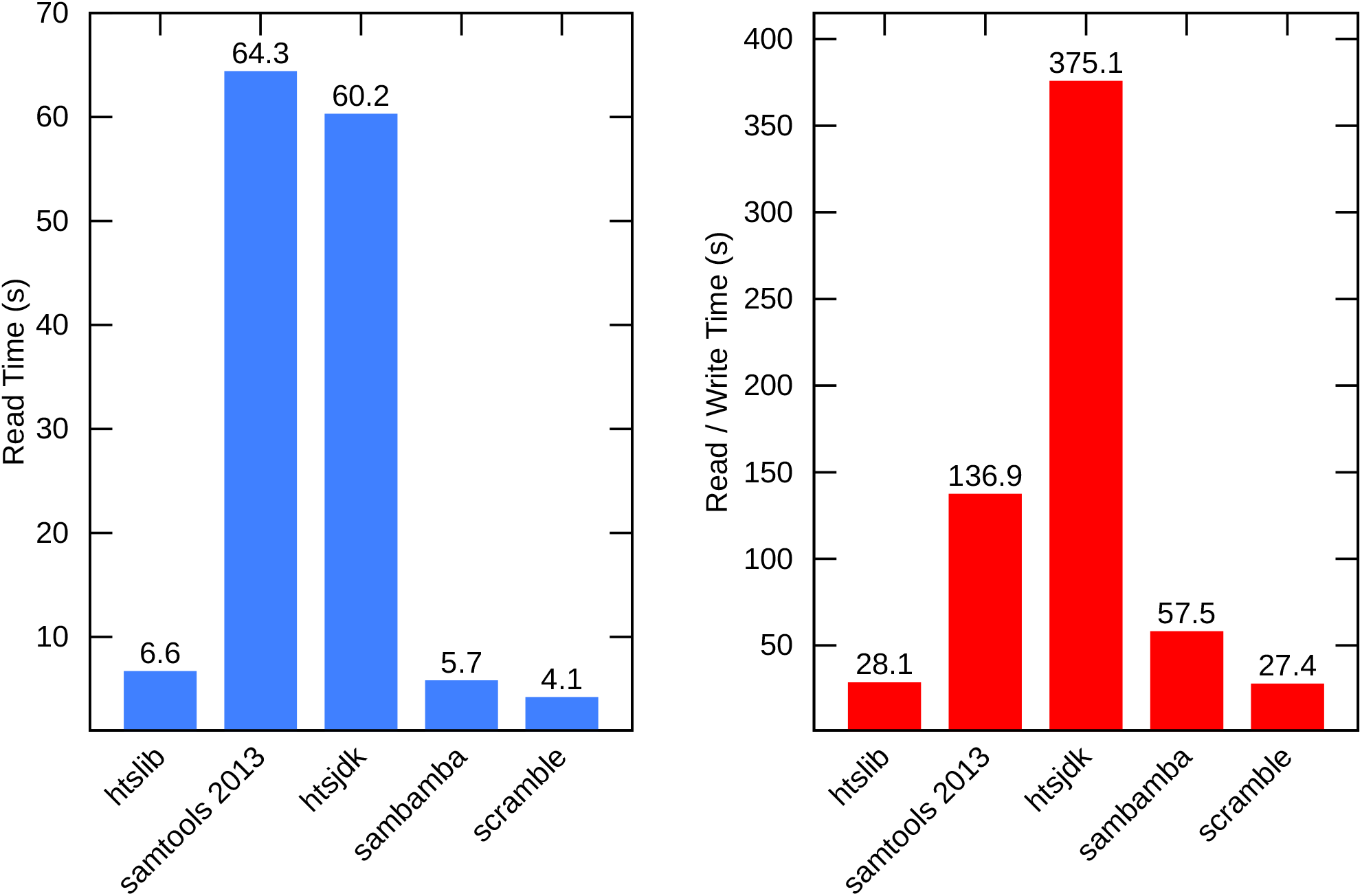
BAM elapsed read/write times, with up to 16 threads available. Read and write elapsed timings for the BAM format on chromosome 1 of ENA accession ERR3239276. Note “samtools 2013” refers to SAMtools version 0.1.19 and not the current SAMtools release. Other tool versions are HTSlib 1.10.2-32-ga22a0af, HTSJDK 2.22.0, Sambamba 0.7,1 and Scramble 1.14.13. These use up to 16 threads, but the HTSJDK times are with only one additional thread per file. SAMtools 0.1.19 has multi-threaded writing only, so the speed is limited by the reading portion. [See Additional file, Section S4 for full single-threaded and multi-threaded timings, along with benchmarks for the SAM and CRAM formats.]

The tests were performed on a RAM disk (/dev/shm) so represent maximum I/O rates for this system.

HTSlib was the only tool tested capable of multi-threaded SAM decoding and encoding, which is important when processing output from a fast multi-threaded aligner. The use of the faster compression library libdeflate[15] over Zlib[16] is also a major contributing factor in BAM performance, meaning BAM to BAM transcoding with 16 threads is 5 times faster than the original SAMtools 0.1.19 and BAM to SAM is 13 times faster.

File sizes also differ slightly for BAM, due to differing Deflate implementations (Zlib, Libdeflate and Intel deflate). HTSlib’s CRAM size is 24% smaller than HTSJDK, while being 4 times faster (with a single thread), although the files remain compatible [see Additional file, Table S5].

To compare the random access capabilities of HTSlib we chose gene and exon regions from the Ensembl database across chromosome 1 and measured the time and I/O statistics to retrieve all alignments overlapping those regions. HTSlib, HTSJDK and Sambamba all support a multi-region iterator that is able to optimise I/O for many regions, reporting alignments that overlap multiple regions once only. SAMtools 0.1.19 and Scramble have no such feature, hence regions that overlap will report some records multiple times and the same block may be read more than once.

The exon list had 58,160 regions (many overlapping each other) covering 5.5% of the chromosome. Figure 3 shows the random access efficiency, in both time and number of bytes read, for the exon list with BAM input. HTSlib is the fastest and requires less I/O to retrieve the same records.

**Figure 3:**
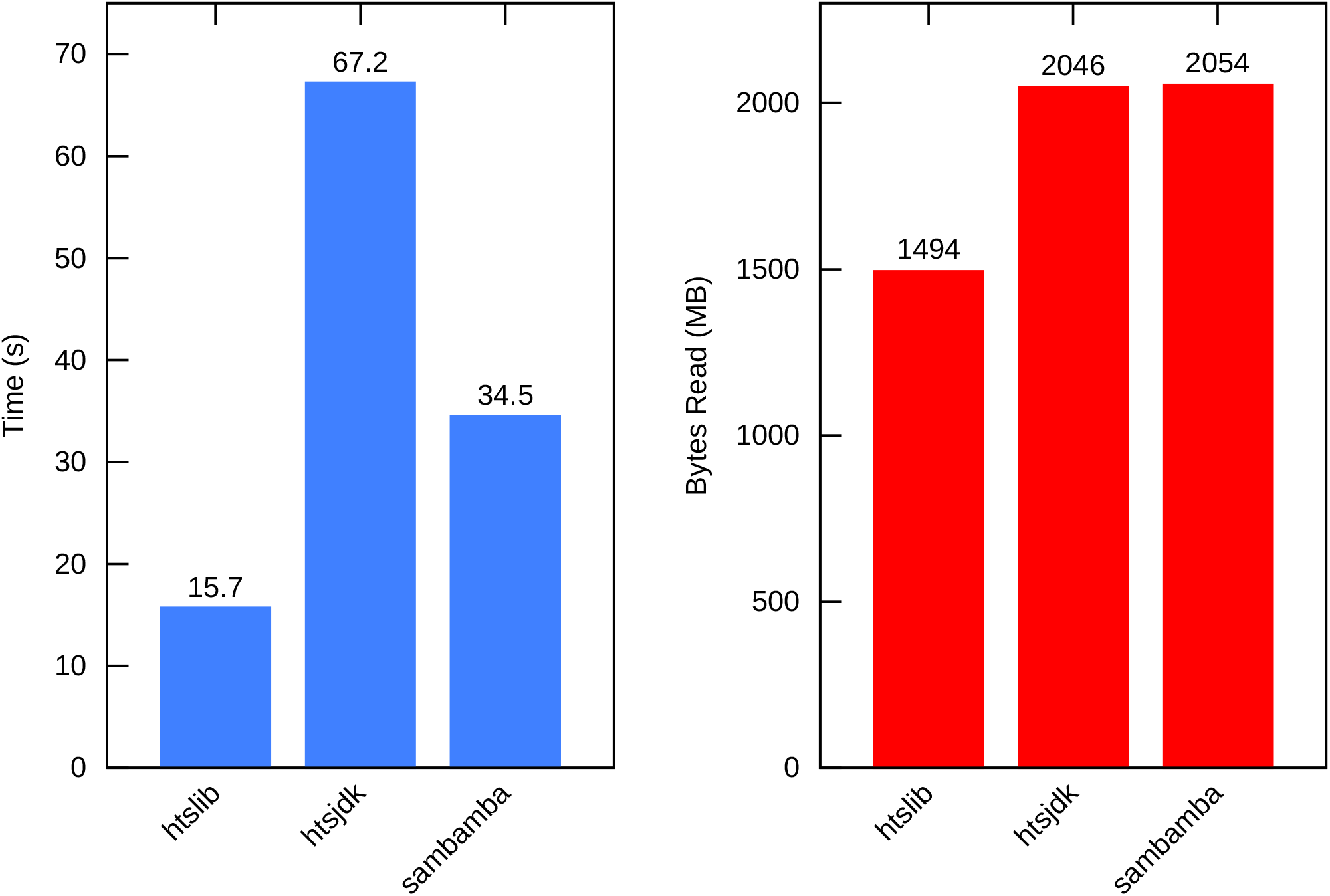
Time and Megabytes of data read, for exon list in BAM. Single threaded performance to read the records overlapping Ensembl exons from chromosome 1.

### Benchmarks of variant formats

The only common format supported between current HTSlib / BCFtools and HTSJDK is BGZF compressed VCF. Figure 4 shows the time to read and read/write this format on a 929 sample test set [17] [see Additional file, Section S8]. Only single thread times are shown as currently multi-threading is sub-optimal in BCFtools and not available HTSJDK.

**Figure 4:**
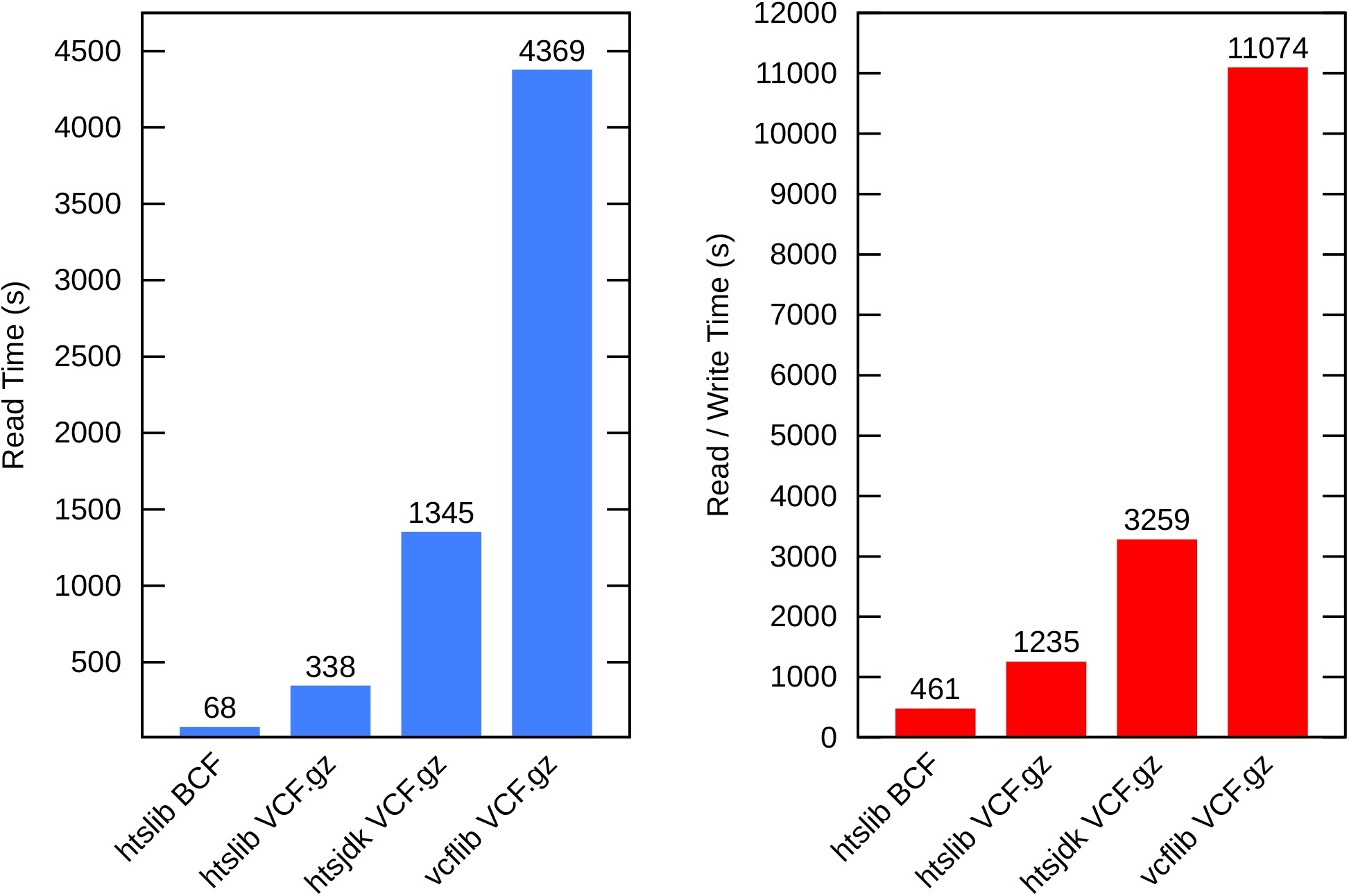
VCF.gz and BCF Read and Read/Write Times, 929 samples. Time in seconds to read and read/write BGZF compressed VCF and BCF. Source file is hgdp_wgs.20190516.full.chr20.vcf.gz, aligned and called from ENA PRJEB6463.

HTSlib also supports the BGZF compressed BCF format, a binary variant of VCF. This is considerably more performant than the compressed VCF, being 5 times faster to decode and nearly 3 times faster to encode. [See Additional file, Tables S10 and S11 for details and more complete results.]

## Discussion

Over the lifetime of HTSlib the cost of sequencing has decreased by approximately 100-fold with a corresponding increase in data volume [18]. New sequencing technologies have also been developed that produce much longer reads. The alignment and variant file formats have moved on from being group led research to being maintained by the File Formats subgroup of the Global Alliance for Genomics and Health[19]. Over the years various improvements and modifications have been made to the specifications. Together these have been and will continue to be a driving force for continued development.

Since HTSlib 1.0, there have been 11 major releases and over 1600 code commits, more than doubling the number of lines of C code. It has gained support for the CRAM file format, better indexing, extended APIs, more transfer protocols (S3, Google Cloud, Htsget) and improved threading and speed. Through the use of automated tests, static analysis tools and fuzz testing it has been made much more reliable [see Additional file, Section S12].

Some of the existing limitations in HTSlib come from the design of the underlying file formats, for example BAM, CRAM and BCF limit the maximum reference length to 2 Gbases [see Additional file, Section S13. We expect future standards development to include improvements leading to better scaling of many-sample VCF; additional support for structural variation; better handling of very long sequencing reads; large genomes; and support for base modifications. Further plans include speeding up both the VCF parser and mpileup, improved documentation, and better support for BED files.

## Availability of supporting source code and requirements

Project name: HTSlib

Project home page: https://www.htslib.org, https://github.com/samtools/htslib

Operating system(s): Platform independent

Programming language: C

License: A mix of Modified 2-Clause BSD (CRAM) and MIT/Expat (everything else).

RRID: SCR_002105

biotools:htslib

## Availability of supporting data

The data set supporting the benchmarking results of this article is available in the European Nucleotide Archive, [ERR3239276 and PRJEB6463], and ftp://ngs.sanger.ac.uk/production/hgdp/hgdp_wgs.20190516/hgdp_wgs.20190516.full.chr20.vcf.gz.

## Additional Files

Additional File, Figure S1: Binary BCF vs VCF format

Additional File, Section S2: Estimated number of HTSlib source code clones

Additional File, Section S3: CRAM Compression Algorithm

Additional File, Section S4: Performance of HTSlib’s SAM, BAM, CRAM implementations

Additional File, Table S5: Read and Read/Write timings for tools and file formats

Additional File, Table S6: Random access times and data volumes, single thread

Additional File, Table S7: Random access times and data volumes, 8 threads

Additional File, Section S8. Performance of HTSlib’s VCF, BCF implementations

Additional File, Figure S9: VCF and BCF read and read/write speeds

Additional File, Table S10: Multi-sample VCF and BCF performance

Additional File, Table S11: Single-sample VCF and BCF performance

Additional File, Section S12: Automatic testing

Additional File, Section S13: The format size limitations

## Abbreviations

API: Application Programming Interface
BAM: Binary sequence Alignment/MAP
BAQ: Base Alignment Quality
BCF: Binary variant Call Format
BGZF: Blocked GNU Zip Format
SAM: Sequence Alignment/Map
VCF: Variant Call Format.

## Authors’ contributions

J.B., P.D., R.D., H.L., J.M., V.O., and A.W. wrote the software. R.D., T.K., and J.M. supervised the project. J.B., P.D., R.D., and A.W. wrote the original draft of the manuscript with all authors reviewing.

## References

1. The 1000 Genomes Project Consortium. A global reference for human genetic variation. Nature. 2015;526: 68–74.

2. Li H, Handsaker B, Wysoker A, Fennell T, Ruan J, Homer N, et al. The Sequence Alignment/Map format and SAMtools. Bioinformatics. 2009;25: 2078–2079.

3. Danecek P, Auton A, Abecasis G, Albers CA, Banks E, DePristo MA, et al. The variant call format and VCFtools. Bioinformatics. 2011;27: 2156–2158.

4. Li H. A statistical framework for SNP calling, mutation discovery, association mapping and population genetical parameter estimation from sequencing data. Bioinformatics. 2011;27: 2987–2993.

5. samtools. HTSJDK. In: GitHub [Internet]. [cited 4 May 2020]. Available: https://github.com/samtools/htsjdk

6. Tarasov A, Vilella AJ, Cuppen E, Nijman IJ, Prins P. Sambamba: fast processing of NGS alignment formats. Bioinformatics. 2015;31: 2032–2034.

7. Bonfield JK. The Scramble conversion tool. Bioinformatics. 2014;30: 2818–2819.

8. Hsi-Yang Fritz M, Leinonen R, Cochrane G, Birney E. Efficient storage of high throughput DNA sequencing data using reference-based compression. Genome Res. 2011;21: 734–740.

9. CRAM. [cited 6 Nov 2020]. Available: https://www.ga4gh.org/cram/

10. Buels R, Yao E, Diesh CM, Hayes RD, Munoz-Torres M, Helt G, et al. JBrowse: a dynamic web platform for genome visualization and analysis. Genome Biology. 2016. doi:10.1186/s13059-016-0924-1

11. Duda J. Asymmetric numeral systems: entropy coding combining speed of Huffman coding with compression rate of arithmetic coding. 2013. Available: https://arxiv.org/abs/1311.2540

12. Kelleher J, Lin M, Albach CH, Birney E, Davies R, Gourtovaia M, et al. htsget: a protocol for securely streaming genomic data. Bioinformatics. 2019;35: 119–121.

13. Li H. Improving SNP discovery by base alignment quality. Bioinformatics. 2011;27: 1157–1158.

14. Li H. Klib. [cited 27 Nov 2020]. Available: https://github.com/attractivechaos/klib

15. Website. [cited 6 Nov 2020]. Available: Eric Biggers, https://github.com/ebiggers/libdeflate

16. Deutsch P, Gailly J-L. ZLIB Compressed Data Format Specification version 3.3. 1996. doi:10.17487/rfc1950

17. Bergstrom A, McCarthy SA, Hui R, Almarri MA, Ayub Q, Danecek P, et al. Insights into human genetic variation and population history from 929 diverse genomes. Science. 2020;367: eaay5012.

18. DNA Sequencing Costs: Data. [cited 23 Sep 2020]. Available: https://www.genome.gov/about-genomics/fact-sheets/DNA-Sequencing-Costs-Data

19. Birney E, Vamathevan J, Goodhand P. Genomics in healthcare: GA4GH looks to 2022. Genomics. bioRxiv; 2017. p. 359.

