## Supplemental Information for "HTSlib - C library for reading/writing high-throughput sequencing data"

**Figure S1. Binary BCF vs VCF format**

BCF, the binary representation of the VCF format, is much faster to process than VCF for two reasons. First, it avoids the expensive conversion from text to the internal binary representation. Second, the fields in BCF are rearranged to allow rapid access to specific value of any sample. This is achieved by storing values in blocks of the same type rather than per-sample, with offsets to blocks determined on the fly.

**VCF**

```
##fileformat=VCFv4.2
##contig=<ID=2,length=51304566>
##INFO=<ID=AC,Number=A,Type=Integer,Description="Allele count in genotypes">
##INFO=<ID=AN,Number=1,Type=Integer,Description="Total number of alleles in called genotypes">
##FORMAT=<ID=GT,Number=1,Type=String,Description="Genotype">
##FORMAT=<ID=DP,Number=1,Type=Integer,Description="Read Depth">
##FORMAT=<ID=GQ,Number=1,Type=Integer,Description="Genotype Quality">
#CHROM  POS ID  REF ALT QUAL FILTER  INFO FORMAT     SAMPLE1     SAMPLE2     SAMPLE3     SAMPLE4     SAMPLE5     SAMPLE6     SAMPLE7
2  81170  .  C  T   .  .    AC=9;AN=7424  GT:DP:GQ   0/0:4:12    0/0:3:9     0/1:1:3     0/1:9:24    1/0:4:12    0/0:5:15    0/0:4:12
2  81171  .  G  A   .  .    AC=6;AN=7446  GT:DP:GQ   0/1:4:12    0/0:3:9     0/0:1:3     0/0:9:24    0/1:4:12    0/1:5:15    0/0:4:12
2  81182  .  A  G   .  .    AC=5;AN=7506  GT:DP:GQ   0/0:5:15    0/0:4:12    0/0:5:15    0/0:9:24    0/0:4:12    0/0:4:12    0/0:4:12
2  81204  .  T  G   .  .    AC=2;AN=7542  GT:DP:GQ   1/0:5:15    0/0:9:27    0/0:10:30   0/0:15:39   0/0:9:27    1/0:13:39   0/1:14:42
```

**BCF**

```
2  81170  .  C  T   .  .    AC=9;AN=7424  GT:0/0:0/0:0/1:0/1:1/0:0/0:0/0    DP:4:3:1:9:4:5:4         GQ:12: 9: 3:24:12:15:12
2  81171  .  G  A   .  .    AC=6;AN=7446  GT:0/1:0/0:0/0:0/0:0/1:0/1:0/0    DP:4:3:1:9:4:5:4         GQ:12: 9: 3:24:12:15:12
2  81182  .  A  G   .  .    AC=5;AN=7506  GT:0/0:0/0:0/0:0/0:0/0:0/0:0/0    DP:5:4:5:9:4:4:4         GQ:15:12:15:24:12:12:12
2  81204  .  T  G   .  .    AC=2;AN=7542  GT:1/0:0/0:0/0:0/0:0/0:1/0:0/1    DP:5:9:10:15:9:13:14     GQ:15:27:30:39:27:39:42
```

**Section S2. Estimated number of HTSlib source code clones**

The number of HTSlib source code clones on github was estimated as follows:

- 352: Number of HTSlib forks on github
  https://github.com/samtools/htslib/network/members

- 908 (est.): Number of C projects linking against HTSlib, whether external or bundled with the project.
  (1853 hits, estimated 49% unique projects)

  *curl -H "Authorization: token $TOK" 'https://api.github.com/search/code?per_page=100&page='$page'&q="-lhts"+AND+htslib"'*

- Number of python projects using pysam .. estimated 9.7k (similar to above)

  *curl -H "Authorization: token $TOK" 'https://api.github.com/search/code?per_page=100&page='$page'&q="import+pysam"'*

**Section S3. CRAM Compression Algorithm**

CRAM stores data in **containers**, with each container having a series of **blocks** which in turn hold one or more **data series**. (A data series is a specific type of data, similar to the columns in SAM.) There is a lot of flexibility in the CRAM specification as to which data series goes into which block, which compression algorithms (codecs) to use on each block, and any parameters for those codecs. CRAM dictates the decoding process, but different CRAM encoders can yield substantially different CRAM efficiency, both in size and speed (see Table S5 below)..

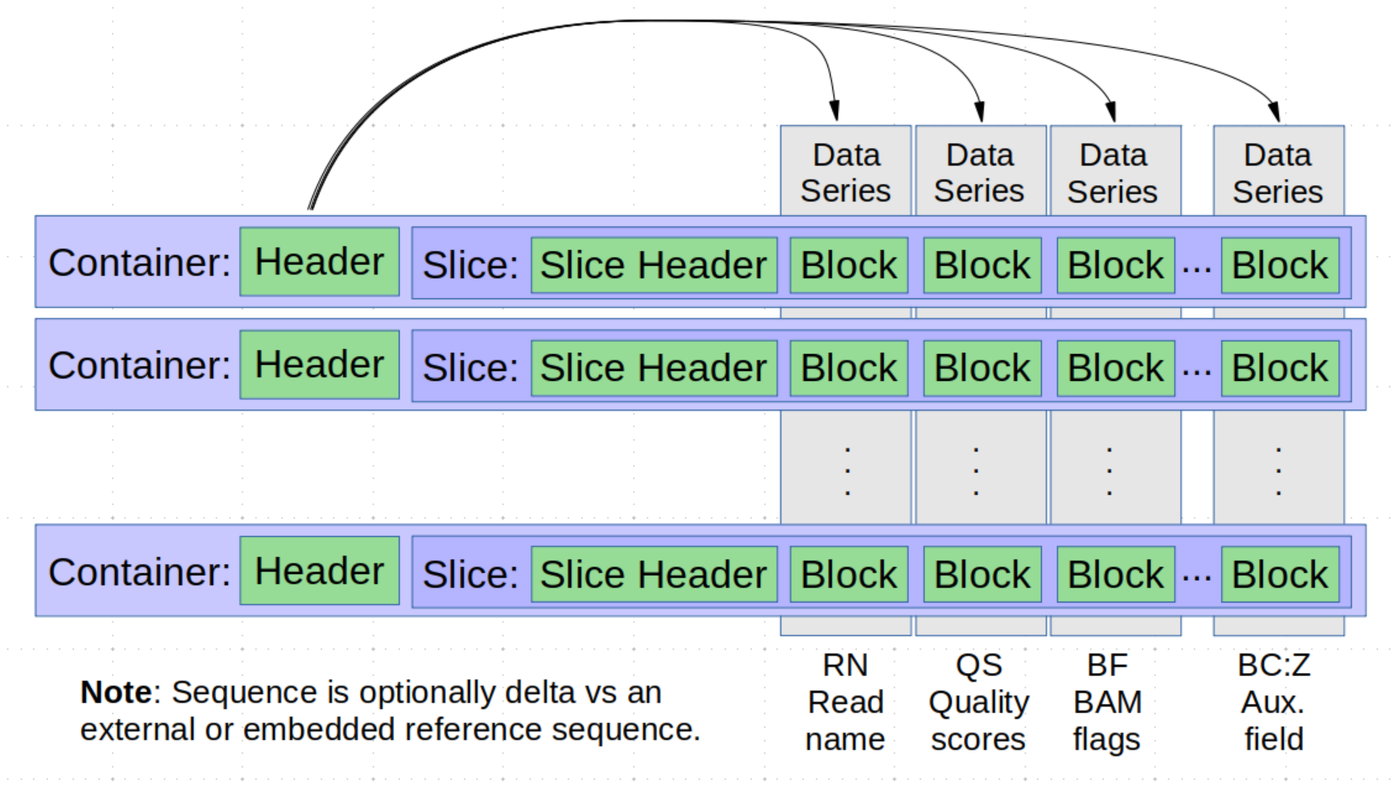

Legend: The structure of a CRAM file. Records are grouped together into containers and slices. Data is then separated into data series (loosely related to a SAM column) and stored in blocks, as described in the container header. Each block is compressed using its own compression codec.

During CRAM encoding, HTSlib initially uses all the codecs it has been permitted to use. With CRAM 3.0 initially this will be Deflate (gzip), Order-0 rANS and Order-1 rANS, however the user may elect to also include bzip2 and LZMA. The draft CRAM 3.1 standard increases the choice of codec further still.

For the first container, each block is compressed with all available codecs and a score recorded, based primarily on the size but also on expected algorithm speed. The weights for speed vs size are adjusted by the compression level, with level 1 favouring speed and level 9 favouring smaller sizes. The best scoring codec is then used to compress that block. This is repeated for the next two containers, aggregating the scores, after which the best overall performing codec is then used for the next 50 containers. This switching between compression trials and usage is repeated, but a method continually failing in the trial period to be within 20% of the best method will be removed from future trials. Finally switching between containers holding mapped data and unmapped data will reset the learnt compression codec behaviour as the style of data may change considerably.

While the CRAM specification permits mixing of multiple data series in a block, HTSlib only stores one data series per block. This offers the best scope for selective random access - both by region but also by type of data. During decode HTSlib can use this to avoid decompressing data if it knows it will never be needed. For example “*samtools flagstat*” does not require the sequence, quality or auxiliary tag data to be decoded.

HTSlib tracks all data series and their dependencies; which data series co-locate in the same block and which data series require others to have previously been decoded (for example insertion sequence needs the feature codes and number of features).

### Section S4. Performance of HTSlib's SAM, BAM, CRAM implementations

Tool versions used:

- HTSlib 1.10.2-32-ga22a0af (https://github.com/samtools/htslib/tree/htslib_paper_benchmarking)
- SAMtools 0.1.19
- HTSJDK 2.22.0
- Sambamba 0.7.1 (https://github.com/biod/sambamba/)
- Scramble 1.14.13
- Vcflib 1.0.1

To time the decode speeds for each tool without consuming additional time performing some complex analysis is not always obvious. Some tools provide built-in benchmark modes, while others need additional tools writing or selection of minimal-processing options. The following method was to benchmark the implementations:

- HTSlib was timed using "*test_view -B*" (benchmark mode).
- SAMtools 0.1.19 was timed using "*samtools view -c*" (count entries).
- HTSJDK required custom java code to use the library. This is available in our git repository. Two variants are available, a non-threaded version and one using asynchronous I/O. This latter version is not fully threaded, but can use up to 3 parallel threads for decode, process and encode.
- Sambamba was timed using "*sambamba view -F supplementary -c*" (count supplementaries). Without the "*-F supplementary*" sambamba counts SAM entries without parsing the data, doing little more counting newlines.
- Scramble was timed using "*scram_flagstat -b*" (benchmark mode).

Scripts to run the benchmarks and produce the tables are in the github repository at https://github.com/jkbonfield/HTSlib_benchmark.

The system used for benchmarking is a virtual machine with 26 cores reported as 2.6Ghz Intel Broadwell, with 214GB of available RAM. The file system for random access tests was a local SSD drive, and for pure CPU throughput tests we used a RAM-disk (/dev/shm). This was necessary to test the maximum theoretical SAM throughput (approximately 5GB/s), which represents the speed at which HTSlib could consume data from a pipe. Throughput is shown below with colour-coding to highlight where a typical HDD and SDD may start to become I/O rate limited.

Table S5 has complete results for HTSlib 1.10, SAMtools 0.1.19, HTSJDK 2.22.0, Sambamba 0.7.1 and Scramble 1.14.13. Formats tested include BAM, SAM (uncompressed), BGZF-compressed SAM, CRAM 3.0 and the draft CRAM 3.1. The latter is not an official GA4GH sanctioned file format and it may still change, but the results indicate the likely performance once ratified.

Benchmarks were tested with no threading enabled (although Java by default multi-threads the garbage collector and this was not disabled), and with 16 threads. Note that HTSJDK does not support multi-threaded decoding and encoding at the time of writing, but it does have an asynchronous mode where one additional thread is used per file descriptor. This mode was enabled for thread tests, but it is only equivalent to a 2 or 3 thread benchmark.

**Table S5**: Read and Read/Write timings for tools and file formats, with and without threading.

| | | No threads, Time and Rate | | | | | 16 Threads, Time and Rate | | | | | |
|---|---|---|---|---|---|---|---|---|---|---|---|---|
| | | Read | | Read/Write | | Index | Read | | Read/Write | | Index | |
| **Tool** | **Format** | **Seconds** | **MB/s** | **Seconds** | **MB/s** | **Seconds** | **Seconds** | **MB/s** | **Seconds** | **MB/s** | **Seconds** | **Size MB** |
| htslib 1.10 | bam | 30.09 | 121.44 | 310.58 | 11.77 | **31.03** | 5.57 | 656.01 | 23.44 | 155.89 | 6.60 | 3654 |
| htslib 1.10 | sam | 39.66 | 848.79 | 94.65 | 355.66 | n / a | 6.23 | 5403.37 | 15.13 | 2224.92 | n / a | 33663 |
| htslib 1.10 | sam.gz | **73.47** | **47.31** | **400.31** | 8.68 | **76.10** | **5.45** | **637.80** | **32.31** | **107.58** | **6.69** | **3476** |
| htslib 1.10 | cram | **73.61** | **17.70** | **304.29** | **4.28** | **0.42** | 11.34 | 114.90 | 43.00 | 30.30 | **0.44** | **1303** |
| htslib 1.10 | cram 3.1 | 61.98 | 17.80 | 292.35 | 3.77 | **0.39** | 11.27 | 97.87 | 43.77 | 25.20 | **0.41** | **1103** |
| samtools 0.1.19 | bam | 51.92 | 69.20 | *549.03* | 6.54 | 52.05 | 52.10 | 68.96 | 113.69 | 31.60 | 52.40 | **3593** |

| | | | | | | | | | | | | |
|---|---|---|---|---|---|---|---|---|---|---|---|---|
| samtools 0.1.19 | sam | 100.10 | 336.29 | 212.26 | 158.59 | n / a | 100.02 | 336.56 | 214.41 | 157.00 | n / a | 33663 |
| htsjdk 2.22.3 | bam | *85.66* | 43.70 | 449.51 | 8.33 | 93.62 | n / a | n / a | n / a | n / a | n / a | 3743 |
| htsjdk | sam | *269.87* | 124.74 | *372.69* | 90.32 | n / a | n / a | n / a | n / a | n / a | n / a | 33663 |
| htsjdk | cram | *297.18* | 5.70 | *1347.04* | 1.26 | n / a | n / a | n / a | n / a | n / a | n / a | *1695* |
| sambamba | bam | 52.22 | 68.81 | 552.92 | 6.50 | 53.60 | 5.12 | 701.76 | 46.64 | 77.04 | **6.54** | 3593 |
| sambamba | sam | 77.62 | 433.69 | 150.19 | 224.14 | n / a | 77.66 | 433.47 | 98.85 | 340.55 | | 33663 |
| sambamba | cram | 87.78 | 16.55 | 335.99 | 4.32 | 0.48 | 26.69 | 54.44 | 56.96 | 25.51 | 0.48 | 1453 |
| scramble | bam | **27.21** | **134.29** | **309.28** | 11.81 | n / a | **3.49** | **1046.99** | **22.73** | **160.76** | n / a | 3654 |
| scramble | sam | **30.89** | 1089.77 | **72.41** | 464.89 | n / a | n / a | n / a | n / a | n / a | n / a | 33663 |
| scramble | cram | 76.14 | 17.14 | 308.63 | 4.23 | 0.49 | **9.75** | 133.85 | **37.12** | **35.16** | 0.49 | 1305 |
| scramble | cram 3.1 | **60.11** | **18.37** | **285.18** | **3.87** | 0.44 | **9.77** | 113.00 | **38.57** | **28.62** | 0.44 | 1104 |

Table S5 - Legend: Single and multi-threaded performance for transcode (read from BAM, write to *format*), decode (read and discard from *format*) and *format* index creation expressed in time (elapsed seconds) and throughput (megabytes per second). To estimate the time to write *format*, subtract the read time from the read/write time for the same *format*. The test data is Chromosome 1 of ENA dataset ERR3685389 (Illumina NovaSeq 6000, Human 40x coverage, sequenced by Illumina, Inc.) in BGZF compressed BAM format. The files were stored on a RAM disk (/dev/shm) to show maximum throughput, but it is acknowledged that storage media may not keep up with the bandwidth. Throughput above 500MB/s (approximately the peak of flashed based SSD) are marked in orange and remaining throughputs between 100MB/s (a typical fast HDD or local network) are marked in blue.

HTSlib has changed considerably since its SAMtools 0.1.19 heritage. The use of libdeflate over zlib means that single threaded BAM decode and encode is considerably faster. The SAM encode and decode has been rewritten and is also over twice as quick. HTSlib's multi-threading has been improved, both efficiency wise but also gaining support for parallel decoding of BAM and encoding/decoding of SAM. These are particularly important when HTSlib is being used to process the SAM output from a rapid multi-threaded alignment tool.

Size wise, there is little difference between the Deflate implementations meaning that BAM sizes are broadly comparable between tools. However HTSlib and Scramble's CRAM implementation offer substantially smaller files than HTSJDK and Sambamba.

Random access was evaluated using the Ensembl GTF file from ftp://ftp.ensembl.org/pub/current_gtf/homo_sapiens/Homo_sapiens.GRCh38.99.gtf.gz.

This contains locations of genes and exons, although the exon list had many overlaps due to alternative splicing. We selected Chromosome 1 only for speed of testing.

The genes and exons BED files were created using the following commands.

```
wget ftp://ftp.ensembl.org/pub/current_gtf/homo_sapiens/Homo_sapiens.GRCh38.99.gtf.gz
zcat Homo_sapiens.GRCh38.99.gtf.gz | awk '$1 == 1 && $3 == "exon" \
        {printf("chr%d\t%d\t%d\n", $1, $4-1, $5)}' > exons.bed
zcat Homo_sapiens.GRCh38.99.gtf.gz | awk '$1 == 1 && $3 == "gene" \
       {printf("chr%d\t%d\t%d\n", $1, $4-1, $5)}' > genes.bed
```

The gene list had 5,475 unique regions and covered around 59% of the chromosome, while the exon list had 58,160 regions covering 5.5% of the chromosome. Such fine scaled random access can be problematic when coupled to coarse granularity CRAM files, so we also tested the efficiency using a CRAM file produced with 1,000 sequences per slice instead of the default 10,000. This grew the CRAM file by 12.6% but in HTSlib this reduced the amount of data to read and time to decode by 52% (although note that the number of seeks increased, which may have an impact when using a traditional disk). Table S6 shows the random access efficiency, in both time and number of bytes read, for the exon list with BAM input.  HTSlib is both faster and also requires less I/O to retrieve the same records.

To demonstrate the impact of the HTSlib multi-region iterator we timed a “samtools view -Mc" vs “samtools view -c” on the BAM file using the exon regions.  With SAMtools-1.10 (using HTSlib) this took 15.5s and 79.3s respectively, yielding 5.0 million and 8.4 million (containing many duplicates) records each.  “SAMtools 0.1.19 view -c” took 142.7s, also giving 8.4 million records.  Counts of the number of reads, seeks and the amount of data read come from the “io_trace” tool (https://github.com/jkbonfield/io_trace).

**Table S6**: Random access times and data volumes, single thread

| | | Exons | | | | | Genes | | | | |
|---|---|---|---|---|---|---|---|---|---|---|---|
| **Tool** | **Format** | **Time** | **#Reads** | **#Seeks** | **MB Read** | **MB/s** | **Time** | **#Reads** | **#Seeks** | **MB Read** | **MB/s** |
| Htslib | BAM | 15.68 | 349642 | 6649 | 1494 | 95 | 22.11 | 554009 | 1760 | 2375 | 107 |
| Htslib | 10k.CRAM | 54.99 | 79919 | 631 | 913 | 17 | 60.31 | 86563 | 360 | 996 | 17 |
| Htslib | 1k.CRAM | 27.56 | 103353 | 7203 | 436 | 16 | 53.92 | 200003 | 2025 | 899 | 17 |
| Htsjdk | BAM | 67.19 | 477007 | 3960 | 2046 | 30 | 69.46 | 597375 | 1992 | 2566 | 37 |
| Htsjdk | 10k.CRAM | 191.74 | 2537813 | 28112 | 1080 | 6 | 214.13 | 2574544 | 28463 | 1097 | 5 |
| Htsjdk | 1k.CRAM | 207.08 | 16283368 | 208596 | 889 | 4 | 393.54 | 19348038 | 245385 | 1064 | 3 |
| Sambamba | BAM | 34.52 | 2156185 | 1681798 | 2054 | 60 | 41.35 | 2694835 | 2098523 | 2570 | 62 |

Table S6 legend:  Evaluation of random access efficiency, in real time and volume of data read, to return records overlapping a target set of regions covering all exons or all genes on chromosome 1.  The 10k.CRAM and 1k.CRAM have 10,000 (default) and 1,000 sequences per slice, demonstrating the effect of improving the granularity of random access in CRAM.  The sizes of these two files were 1303MB and 1467MB respectively.  Bold entries show the smallest value for that format and metric category.  All tests used 1 main thread only.  The files were stored on a local SSD with the file system cache purged between tests.  Throughput above 100MB/s (a typical fast HDD or local network) are marked in blue.

Sambamba had many seeks, but about half of these were `lseek(7, 0, SEEK_CUR)`, i.e. seeking to the current location.

We also tested multi-threading in HTSlib and Sambamba when used with random access.  This is a challenging operation, particularly when reading from many small disparate regions as in the exons.bed file.  The results of 8 threads are shown in Table S7.  Sambamba gains the most here, but not enough to surpass the performance of HTSlib.  Neither tool was able to achieve an 8-fold performance gain.

**Table S7**: Random access times and data volumes, 8 threads

| | | Exons | | | | | Genes | | | | |
|---|---|---|---|---|---|---|---|---|---|---|---|
| **Tool** | **Format** | **Time** | **#Reads** | **#Seeks** | **MB Read** | **MB/s** | **Time** | **#Reads** | **#Seeks** | **MB Read** | **MB/s** |
| Htslib | BAM | 6.32 | 388608 | 6978 | 1660 | 263 | 6.11 | 563411 | 1775 | 2416 | 395 |
| Htslib | 10k.CRAM | 18.1 | 79922 | 631 | 913 | 50 | 15.91 | 86564 | 360 | 996 | 63 |
| Htslib | 1k.CRAM | 17.69 | 103373 | 7203 | 436 | 25 | 13.59 | 200010 | 2025 | 899 | 66 |
| Sambamba | BAM | 8.11 | 2156357 | 1681910 | 2055 | 253 | 7.98 | 2695007 | 2098635 | 2571 | 322 |

Table S7 legend:  Random access performance when given 8 threads.  HTSlib speeds vary between 2-4 times faster than a single thread.  Sambamba speeds are 4-5 times faster.  Throughputs above 100MB/s (a typical fast HDD or local network) are marked in blue.

**Section S8. Performance of HTSlib's VCF, BCF implementations**

Tool versions used:

- HTSlib cram_codecs branch a22a0af *(https://github.com/samtools/HTSlib/pull/929)*
- HTSJDK 2.22.0
- Vcflib 1.0.1

The input data was ftp://ngs.sanger.ac.uk/production/hgdp/hgdp_wgs.20190516/hgdp_wgs.20190516.full.chr20.vcf.gz. This contains 929 merged samples, so we evaluate performance on the full data set and only the first sample from this file. File sizes were 8239MB for the full VCF.gz and 143.4MB for a single sample. In BAM these sizes grew to 9093MB and 152.9MB respectively.

The decoding and encoding time on the multi-sample file is dominated by the VCF FORMAT columns. A pull-request (https://github.com/samtools/htslib/pull/1081) to HTSlib permits decoding the format field to be delayed until required, retaining it in textual form. This greatly decreases the time with the difference being visible in the light shading in Figure S9 and the “Fast-VCF” table column. Note many operations still require access to this data, so the speed will depend on the command being used.

Measurements were made using BCFtools 1.10 linked against the HTSlib pull-request listed above. This optionally adds a faster VCF parsing mode where the FORMAT fields are read into memory and may be written back out again, but are only decoded once necessary such as conversion to BCF or filtering by a FORMAT column. This partial decode strategy is shown in the “Fast-VCF” column of the Tables S10 and S11 below.

None of this data was phased, so we tested two no-op filters with "*bcftools view --min-alleles 0*" (using non-FORMAT columns) and “*bcftools view --exclude-phased*” (using FORMAT columns). Both of these return all data, with the latter required decoding the FORMAT columns. This demonstrates the difference between Fast-VCF.gz and the full VCF.gz columns. To perform read-only tests, these two commands can be negated ("*--min-alleles 999*" and *--phased*) to filter out 100% of records to exclude the cost of writing.

Similarly for Vcflib the *vcffilter* tool was used to select all reads ("*-f 'AN < 999999'*") and no reads ("*-f 'AN > 999999'*"). The difference between using *vcffilter* to report all reads and vcfecho (a simple read/write) loop was around 5%. This cpu time was subtracted from the "*vcffilter -f 'AN > 999999'*" command in order to get a more accurate read-only benchmark.

**Figure S9**: VCF and BCF read and read/write speeds for the 929 sample dataset.

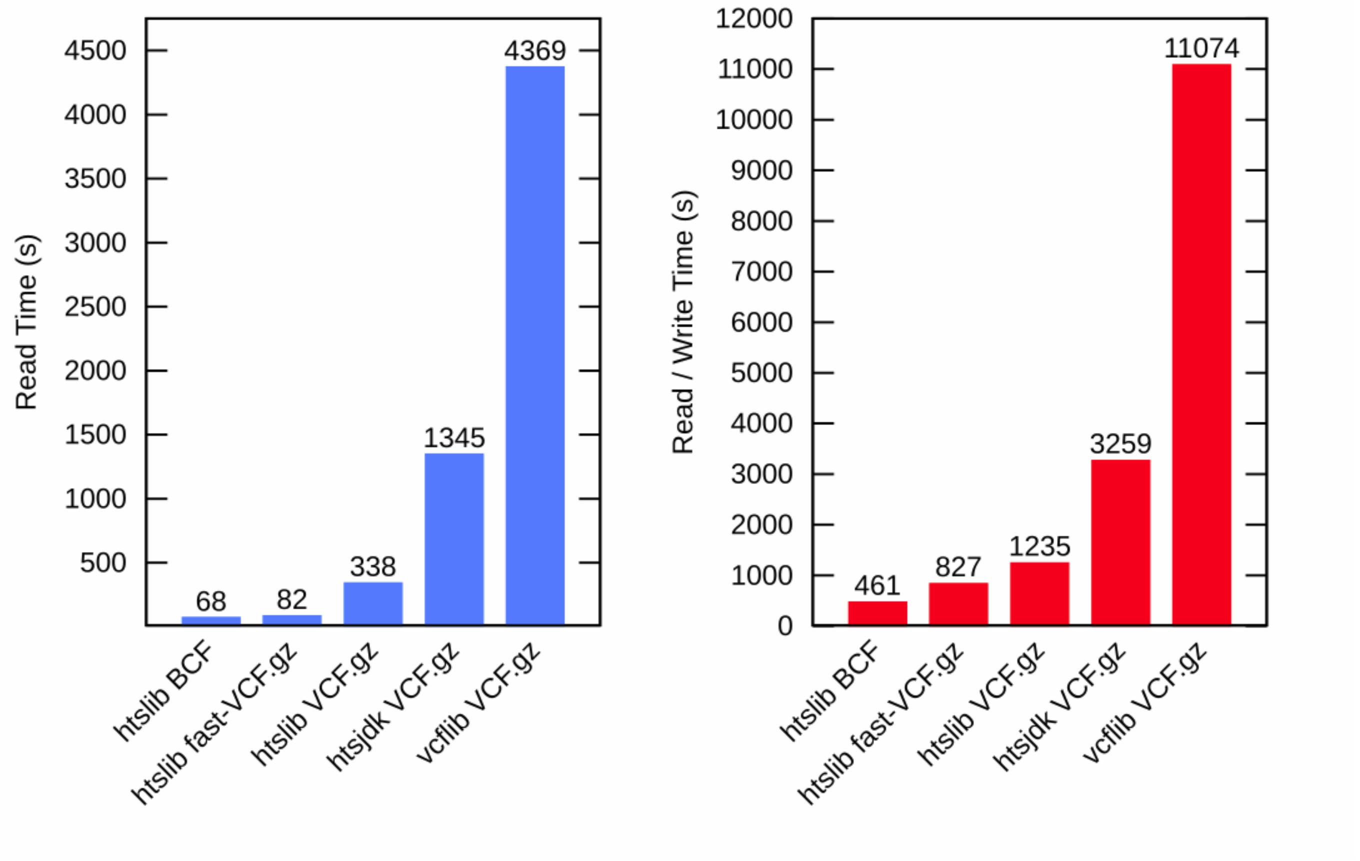

**Table S10**: Multi-sample VCF and BCF performance

| | Fast-VCF.gz | | | | VCF.gz | | | | BCF | | | |
|---|---|---|---|---|---|---|---|---|---|---|---|---|
| | Read | | Read/Write | | Read | | Read/Write | | Read | | Read/Write | |
| **929 samples** | Seconds | MB/s | Seconds | MB/s | Seconds | MB/s | Seconds | MB/s | Seconds | MB/s | Seconds | MB/s |
| **htslib 1.10** | 81.7 | 100.8 | 827.2 | 10.0 | 338.4 | 24.3 | 1234.7 | 6.7 | 66.57 | 136.6 | 461.48 | 19.7 |
| **htsjdk 2.22.0** | | | | | 1344.8 | 6.1 | 3258.9 | 2.5 | | | | |
| **Vcflib 1.0.1** | | | | | 4369.28 | 1.9 | 11073.97 | 0.7 | | | | |

Throughputs above 100MB/s (a typical fast HDD or local network) are marked in blue.

**Table S11**: Single-sample VCF and BCF performance

| | Fast-VCF.gz | | | | VCF.gz | | | | BCF | | | |
|---|---|---|---|---|---|---|---|---|---|---|---|---|
| | Read | | Read/Write | | Read | | Read/Write | | Read | | Read/Write | |
| **1 sample** | Seconds | MB/s | Seconds | MB/s | Seconds | MB/s | Seconds | MB/s | Seconds | MB/s | Seconds | MB/s |
| **htslib 1.10** | 5.33 | 26.9 | 19.38 | 7.4 | 6.7 | 21.4 | 20.94 | 6.8 | 1.72 | 88.9 | 9.41 | 16.2 |
| **htsjdk 2.22.0** | | | | | 14.14 | 10.1 | 40.2 | 3.6 | | | | |
| **Vcflib 1.0.1** | | | | | 25.43 | 5.6 | 132.54 | 1.1 | | | | |

As expected the impact of partial decoding is huge for the multi-sample VCF.  However in all cases BCF is considerably faster, so it is unfortunate that it has not been adopted as the standard interchange format by tool authors.

It is also clear there is a very significant difference between single-sample and multi-sample speeds even when not fully decoding the FORMAT columns.  This is due to the overhead in compression (Deflate) as the multi-sample file is 57 times larger than the single-sample one.  New file formats for many-sample datasets are desirable.

The growth in CPU time moving from 1 sample to 929 samples is not the same ratio for all tools. HTSlib is 50 times slower to fully decode the whole data set than a single sample, while HTSJDK is 95 times slower and vcflib 170 times slower. However with only two data points we cannot say how these trends scale.

### Section S12. Automatic testing

With the widespread use of HTSlib, it is important that the library is reliable. To ensure this, it includes a comprehensive test harness. Continuous integration services run all of the tests on a variety of operating systems (including Linux, MacOS and Windows) and platforms (x86-64 and s390x) whenever code is checked into the source repository, ensuring bugs are caught and fixed rapidly. Code quality is also assured by checking for memory errors using Valgrind memcheck [10] and AddressSanitizer [11]. Additionally, UndefinedBehaviourSanitizer is used to detect violations of the C standard.

As HTSlib may need to read inputs from unknown sources, care has been taken to ensure it behaves in a safe manner even on badly malformed data. It has been subjected to extensive fuzz testing using American Fuzzy Lop [12] and libfuzzer [13]. It has also been included in the Google OSS-Fuzz project, which allows it to be tested with a number of different fuzzers (including those already listed) at very large scale.

#### *Numbers of tests per package*

*HTSlib: 199*

- *11 test_* dedicated programs*
- *13 tests from tabix*
- *21 tests from pileup*
- *154 tests in test.pl*

*Samtools: 702*

- *4 test_* programs*
- *698 tests in test.pl*

*Bcftools: 1426*

- *4 test_rbuf*
- *1 test_regidx*
- *1421 test.pl*

*Total: 2327*

**Section S13. The format size limitations**

- The maximum length of chromosomes that BAI and TBI indices can hold is 512 Mbases.
- The maximum indexable file size for BGZF compressed files is $2^{48}$ bytes.
- The maximum reference size for BAM, CRAM and BCF files is $2^{31}$ bases.
- The BAM and CRAM header sizes are limited to 2 Gbytes.
- The BCF header size is limited to 4 Gbytes.
